# Revisiting Decompression Sickness Risk and Mobility in the Context of the SmartSuit, a Hybrid Planetary Spacesuit

**DOI:** 10.1101/2021.03.26.437246

**Authors:** Logan Kluis, Ana Diaz-Artiles

## Abstract

Gas pressurized spacesuits are cumbersome, cause injuries, and make completing tasks efficiently difficult. Decreasing the gas pressure of the spacesuit is an effective method of improving mobility, but reduction in the total spacesuit pressure also results in a higher risk for decompression sickness (DCS). The risk of DCS is currently mitigated by breathing pure oxygen before the Extravehicular Activity (EVA) for up to 4 hours to remove inert gases from body tissues, but this has a negative operational impact due to the time needed to perform the prebreathe. In this paper, we review and quantify these important trade-offs between spacesuit pressure, mobility, and prebreathe time (or risk of DCS) in the context of future planetary EVAs. These trade-offs are highly dependent on the atmospheric conditions used in the space habitat or space station, and therefore, these conditions are also important considerations for future planetary exploration activities. In our analysis, we include three habitat scenarios (International Space Station: 14.7 psia, 21% O_2_, Adjusted Space Shuttle: 10.2 psia, 26.5% O_2_, and Exploration: 8.2 psia, 34% O_2_) to further quantify these differences. In addition, we explore these trade-offs in the context of the SmartSuit spacesuit architecture, a hybrid spacesuit with a soft robotic layer that, not only increases mobility with assistive actuators in the lower body, but it also applies 1 psia of mechanical counterpressure (MCP). The additional MCP in hybrid spacesuits can be used to supplement the gas pressure (i.e., increasing the total spacesuit pressure), therefore reducing the risk of DCS (or reduce prebreathe time). Alternatively, the MCP can be used to reduce the gas pressure (i.e., maintaining the same total spacesuit pressure), therefore increasing mobility. Finally, we propose a variable pressure concept of operations for the SmartSuit spacesuit architecture, where these two MCP applications are effectively combined during the same EVA to maximize the benefits of both configurations. Our framework quantifies critical spacesuit and habitat trade-offs for future planetary exploration, and contributes to the assessment of human health and performance during future planetary EVAs.

## 1. Introduction

The current United States (US) spacesuit used for spacewalks on the International Space Station (ISS) is the extravehicular mobility unit (EMU)^1,2^. The EMU operates in a microgravity environment at a gas pressure of 4.3 psia (29.6 kPa) and 100% oxygen^3^. The EMU components come in discrete sizes to accommodate a wider range of astronauts but has consequently led to spacesuits that do not optimally fit the entire astronaut population^4–7^. The combination of poor fit and gas pressurization creates a suit environment that is difficult to move in^2,6,8,9^, causes injuries^7,10–14^, and limits range of motion during extravehicular activity (EVA)^15–20^. The next generation spacesuit built for planetary exploration is the xEMU, which is capable of pressurizing up to 8.2 psia (56.5 kPa)^21^. While the xEMU has been designed to ensure better mobility, operations at higher suit gas pressures might intensify some of the existing problems with the EMU, which may lead to suboptimal EVA performance and impact mission success.

A simple solution to impaired mobility due to high gas pressures consists in decreasing the operating pressure of the spacesuit. While lower spacesuit pressures are a viable answer, a trade-off exists between decreasing spacesuit pressure and increasing the risk of decompression sickness (DCS)^22,23^. DCS is characterized by the formation of inert gas bubbles (typically nitrogen) in human tissue due to rapid decompression such as a diver ascending in water or an astronaut entering a lower pressure spacesuit^24–26^. Symptoms of DCS range from pain in the muscles and joints to circulatory collapse, shock, and even death^27^. As a result, DCS is a major risk that must be mitigated to ensure astronaut safety. To address the risk of DCS, NASA’s protocol on the ISS calls for four hours of breathing pure oxygen before an EVA to purge the tissues of nitrogen^28–31^. For a Martian mission that will potentially have almost daily EVAs^32^, it is not operationally practical to require four hours of prebreathe time per EVA.

The risk of DCS is defined as the ratio of nitrogen in the tissue to the pressure in the spacesuit. Oxygen prebreathe reduces the amount of nitrogen in the tissue and thus the risk of DCS, but two other solutions exist: 1) reducing the partial pressure of the nitrogen in the habitat atmosphere, or 2) increasing the pressure of the spacesuit. Decreasing the partial pressure of nitrogen in the habitat by increasing the percentage of oxygen can be costly, increases flammability, and also increases the probability of hyperoxia. Conversely, reducing the partial pressure of oxygen increases the risk of hypoxia. Cabin atmospheres other than the nominal Earthen sea-level atmosphere used on the ISS (14.7 psia and 21% oxygen) have been implemented in space. For example, the Mercury, Gemini, and Apollo missions used pure oxygen at low pressures to avoid DCS and hyperoxia^22^. The Space Shuttle creatively altered the cabin atmosphere before EVA missions to 10.2 psia and 26.5% oxygen to effectively lower the oxygen prebreathe time to 40 minutes^33^. For future missions, the Exploration Atmosphere Working Group (EAWG) examined the trade-offs between DCS, hypoxia, flammability, and several other factors, and recommended an exploration atmosphere of 8.0 psia and 32% oxygen, which nearly eliminates the need to prebreathe^23^. This atmosphere was later increased to 8.2 psia and 34% as it supplies physiological relief without a negative impact to operational capabilities^34^.

In this context, we consider the development of a novel spacesuit architecture for EVA operations on planetary surfaces called SmartSuit. The SmartSuit, while still gas pressurized, incorporates a full-body soft-robotic layer that increases astronaut mobility, therefore decreasing metabolic expenditure facilitating exploration operations^35^. In addition to the enhanced mobility, the soft robotic layer is capable of applying a certain amount of mechanical counter-pressure (MCP). The addition of MCP could reduce the amount of gas pressure needed (which improves the mobility of the spacesuit); or, when combined with the original gas pressure, the MCP could increase the total pressure (which in turn decreases prebreathe time). In this paper, we explore and further analyze the trade-off between spacesuit pressure, risk of DCS, and mobility in the context of the SmartSuit. In particular, and based on our previous investigations on viable SmartSuit architectures, we consider the use case where the SmartSuit soft-robotic actuators are capable of providing up to 10 Nm of assistive torque in the lower body joints (hip, knee, and ankle joints). In addition, we assume that the soft-robotic layer is capable of providing up to 1 psia of MCP. However, our framework provided herein also permits to quickly analyze other hybrid spacesuit architectures and visualize important trade-offs for spacesuit design and operations.

## 2. Factors Under Consideration

### Risk of Decompression Sickness

To quantify the risk of decompression sickness, it is common to use the ratio between the partial pressure of nitrogen in the tissue and the pressure of the spacesuit (also known as the bends ratio)^22^:

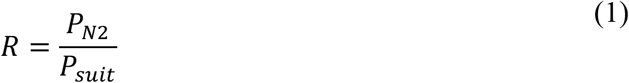

where *P_N2_* is the initial absorbed tissue N_2_ pressure (i.e., cabin N_2_ partial pressure), and *P_suit_* is the total spacesuit pressure. During pure oxygen prebreathe, the elimination of nitrogen follows an exponential decay curve with a tissue dependent half time, *t_1/2_* (typically equal to 360min), that can be expressed in terms of R value^22,36^:

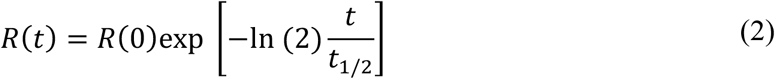

The protocol on the ISS and the Space Shuttle requires a final R in the range of 1.60 and 1.70^22,28^. As an example, using the equations (1) and (2) in the context of the protocol currently in place on the ISS (i.e., R(0) = 2.73 (11.76 psia *P_N2_* and 4.3 psia *P_suit_*), t_1/2_ = 360 min, and t = 240 min (4-hour prebreathe time)), the DCS risk value becomes R = 1.7. Similarly, the final R value for astronauts on the Space Shuttle is calculated using the adjusted Space Shuttle atmosphere (7.5 psia *P_N2_*) and the 40-minute prebreathe time (t = 40min). Thus, using the same spacesuit pressure and tissue dependent half time as the ISS scenario (4.3 psia *P_suit_* and t_1/2_ = 360 min), the DCS risk value is R = 1.61. The actual R values are typically lower as a result of the conservative choice of the tissue dependent half time, *t_1/2_*^22^. For future planetary exploration missions, Conkin recommends a DCS risk value R between 1.3 and 1.4^37^.

### Mobility Scores

Due to gas pressurization, spacesuits constrain mobility and increase the metabolic cost of movement. Our group has performed biomechanical analyses using OpenSim and have quantified the impact that the EMU has on metabolic rate^35,38^, using joint torques obtained from experimental spacesuit testing^2,39,40^. Simulations included a walking motion in which external EMU joint torques were applied to the hips, knees, and ankles. In addition, a second set of simulations were performed with the same external EMU joints combined with additional soft robotic actuators in the hip, knee, and ankle joints that are capable of applying up to 10 Nm of assistive torque to improve joint motion and thus metabolic cost. If we assume that the effect of gas pressure on spacesuit joint torques is linear^41^, we can replicate the simulations for scenarios of reduced gas pressure (for example because the presence of MCP allows to do so), with and without assistive soft robotic actuators. For example, if the pressure in the EMU is decreased by half, the associated external joint torques that the spacesuit wearer needs to counteract while moving inside the spacesuit are also reduced by half. Thus, using the metabolic model developed by Umberger^42,43^and the methodology described in our previous publications^35,38^, we calculated walking energy expenditure in different spacesuit pressure conditions. For our analysis, we focused on the SmartSuit scenario in which the soft-robotic actuation can produce up to 10 Nm of assistive torque, which is consistent with previous prototype testing^38^. The simulated walking conditions are the following: a) only spacesuit joint torques at pressures of 0 psia (unsuited), 1.075 psia (25% of EMU operating pressure), 2.15 psia (50% of EMU operating pressure), 3.225 psia (75% of EMU operating pressure), and 4.3 psia (EMU operating pressure), and b) spacesuit joint torques combined with assistive actuators on the hips, knees, and ankles that are capable of producing up to 10 Nm of torque at the same pressures as a). The walking energy expenditure results from the simulations in the different conditions are summarized in Table 1.

**Table 1:** Walking Energy Expenditure from biomechanical simulations of EMU-suited walking motion, with and without soft-robotic actuators (up to 10N of assistive torque). EMU and robotic joint torques incorporated in the simulation include hips, knees, and ankles^2,39^. Results include the following conditions: unsuited, EMU pressurized at 1.075 psia (25% of EMU operating pressure), EMU pressurized at 2.15 psia (50% of EMU operating pressure), EMU pressurized at 3.225 psia (75% of EMU operating pressure), and EMU pressurized at 4.3 psia (nominal EMU operating pressure). Every condition was simulated with and without robotic torque actuators. EMU joint torques were assumed to scale linearly with gas pressure^41^. Energy expenditure is measured in kcal/hour.

| Scenario | Walking Energy Expenditure:<br>only Spacesuit Joint Torques (kcal/hour) | Walking Energy Expenditure:<br>Spacesuit Joint Torques and Assistive Robotic Actuators (kcal/hour) |
| --- | --- | --- |
| Unsuited | 510 | 329 |
| EMU pressurized at 1.075 psia (25% of EMU operating pressure) | 661 | 548 |
| EMU pressurized at 2.15 psia (50% of EMU operating pressure) | 705 | 600 |
| EMU pressurized at 3.225 psia (75% of EMU operating pressure) | 865 | 753 |
| EMU pressurized at 4.3 psia (nominal EMU operating pressure) | 942 | 799 |

For our subsequent analysis, we focus on a SmartSuit scenario in which the soft-robotic actuation layer can also produce between 0 and 1 psia of MCP. Because the MCP is applied from a skin tight, soft robotic layer, we assume there will be no additional penalty to mobility for replacing gas pressure with any amount of MCP. Based on results from Table 1, the relationship between the amount of gas pressure and the metabolic cost appears to be approximately linear. Thus, we define a mobility score that we derived from the energy expenditure simulations with joint torque actuators shown in Table 1 using a linear fit model. The scores were then normalized by the energy expenditure calculated for unsuited walking without the robotic actuators (510 kcal/hour). As a result, a mobility score of 2 represents a spacesuit scenario in which energy expenditure during ambulation is twice as expensive as that of unsuited walking. Figure 1 presents the relationship between mobility score and gas pressure.

**Figure 1:**
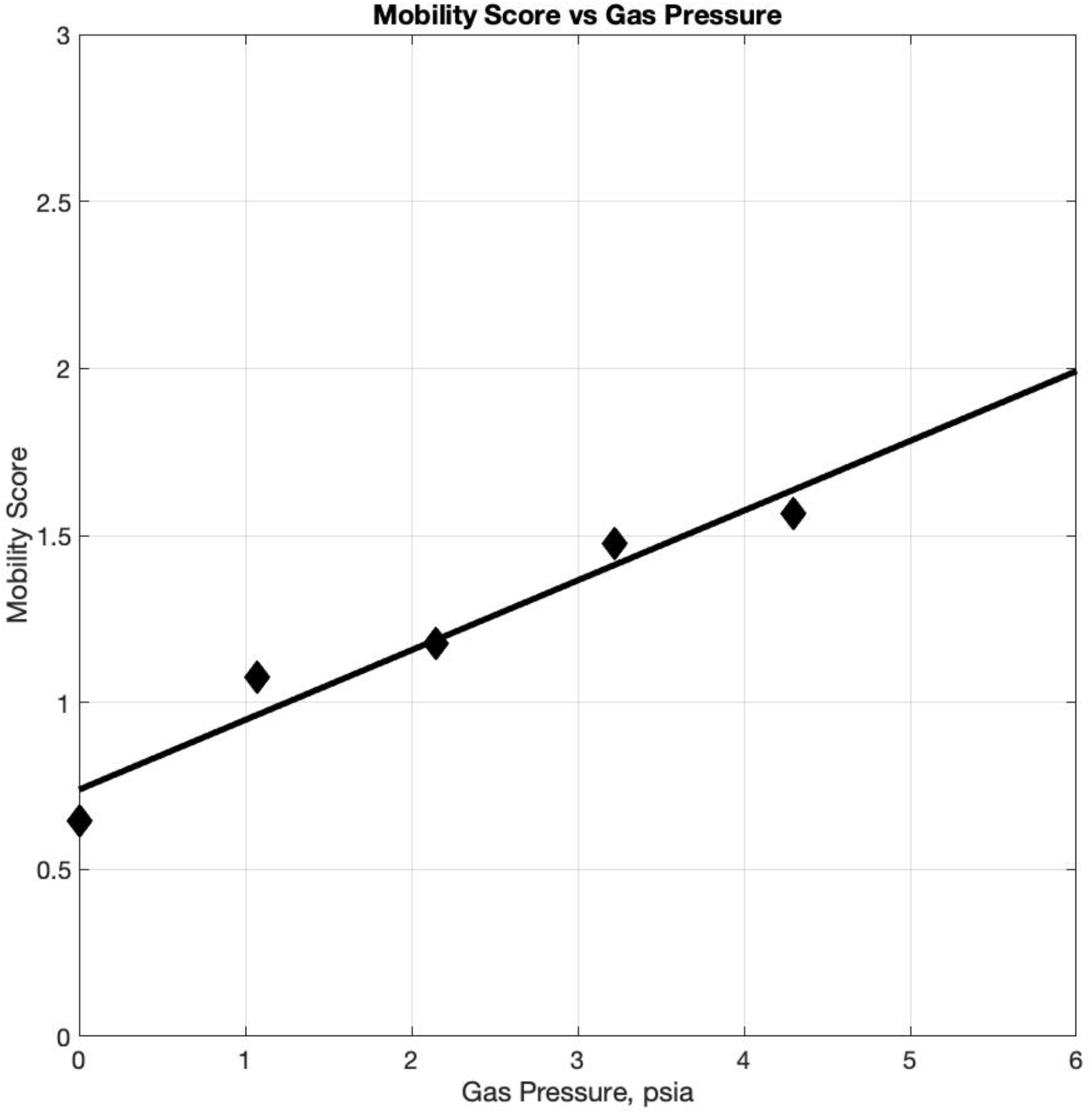
Mobility score as a function of gas pressure in the spacesuit. The mobility score is derived from the energy expenditure simulations with assistive robotic actuators shown in Table 1. These scores are then normalized by the metabolic cost of unsuited walking without actuators (510 kcal/hour). As a result, a mobility score of 2 represents a spacesuit environment that requires twice the amount of energy expenditure as that of unsuited walking.

### Atmospheres

The cabin atmosphere has a critical role in the determination of DCS risk and thus, prebreathe time. The EAWG was tasked with recommending a habitat atmosphere for future planetary missions. To do so, they considered four risks: hypoxia, flammability, mission impact, and DCS^23^. Hypoxia refers to the condition of breathing lower partial-pressure oxygen for an extended period of time. Side effects include decrements in vision^44,45^, cognitive performance^46^, acute mountain sickness^47,48^, and overall reduction in mission performance^49^. Flammability refers to the risk of materials to catch fire in the presence of different oxygen levels. As the percent of oxygen in the air increases, so does the flammability^50^. Typically, the testing limit for general materials in a spacecraft habitat is set to an oxygen limit 30%^22^but, according to the EAWG, a 36% oxygen atmosphere is supportable with the materials available^23^. The third risk considered refers to mission impact, specifically related to astronaut performance decrements during EVAs when operating in a highly pressurized environment, which could potentially impact mission success. The fourth and final risk considered refers to DCS, which has already been described in previous paragraphs. After considering these risks, the EAWG recommended an atmosphere of 8.0 psia and 32% oxygen for future planetary missions. This atmosphere was later changed to 8.2 psia and 34% oxygen as an increase in pressure and oxygen content lowered the risk of hypoxia while remaining in a suitable range for flammability risk^34^. Thus, our analysis includes these atmospheric conditions that we refer to as Exploration atmosphere. In addition, we also include the current ISS cabin atmosphere of 14.7 psia and 21% oxygen, which is an Earth-normal habitat environment. Finally, the Space Shuttle functioned at an identical atmosphere than the ISS but also had the capability to operate at a lower pressure and higher oxygen environment (10.2 psia and 26.5% oxygen) before an EVA to reduce prebreathe time. This Adjusted Space Shuttle environment was also included in our analysis. A summary of the atmospheres considered can be found in Table 2.

**Table 2:** The habitat atmospheric conditions (pressure and oxygen concentration) considered in the present analysis.

| Habitat Atmospheric Conditions | Cabin Pressure, psia | Cabin Oxygen Concentration, % |
| --- | --- | --- |
| ISS | 14.7 | 21 |
| Adjusted Space Shuttle | 10.2 | 26.5 |
| Exploration | 8.2 | 34 |

## 3 Trade-off Analysis

### Risk of Decompression Sickness vs. Prebreathe Time vs. Spacesuit Pressure

We conducted a trade space analysis between the risk of DCS, prebreathe time, and operational spacesuit pressure. The risk of DCS is calculated using equations 1 and 2 for a given spacesuit pressure, cabin atmosphere, and prebreathe time. Using the previous examples, a 4-hour (i.e., 240 min) pure oxygen prebreathe time before donning a 4.3 psia spacesuit on the ISS results in a risk factor of approximately R = 1.7, as shown in Figure 2, top panel. Similarly, the protocol for the Space Shuttle required a 40-min prebreathe (after 36 hours at the lower pressure environment of 10.2 psia and 26.5% oxygen) before donning the EMU (pressurized at 4.3 psia) to maintain a risk factor between 1.6 and 1.7 (see Figure 2, middle panel). In the case of the Exploration atmosphere, a prebreathe of 240 minutes returns a DCS risk of R = 0.8 (for a spacesuit of 4.3 psia), while a 40-min prebreathe for the same spacesuit pressure yields a DCS risk of approximately R = 1.2 (see Figure 2, bottom panel). If the spacesuit is pressurized to 8.3 psia instead, the risk of DCS after a 40-min pure oxygen prebreathe decreases to R = 0.6. Figure 2 shows the spacesuit design space considered and quantifies the existing trade-offs for multiple habitat atmospheres. It also demonstrates the benefits of using Exploration-type atmospheres from the point of view of DCS risk and prebreathe time, especially if higher spacesuit pressures (e.g., 8.2 psia for the xEMU) are considered.

**Figure 2:**
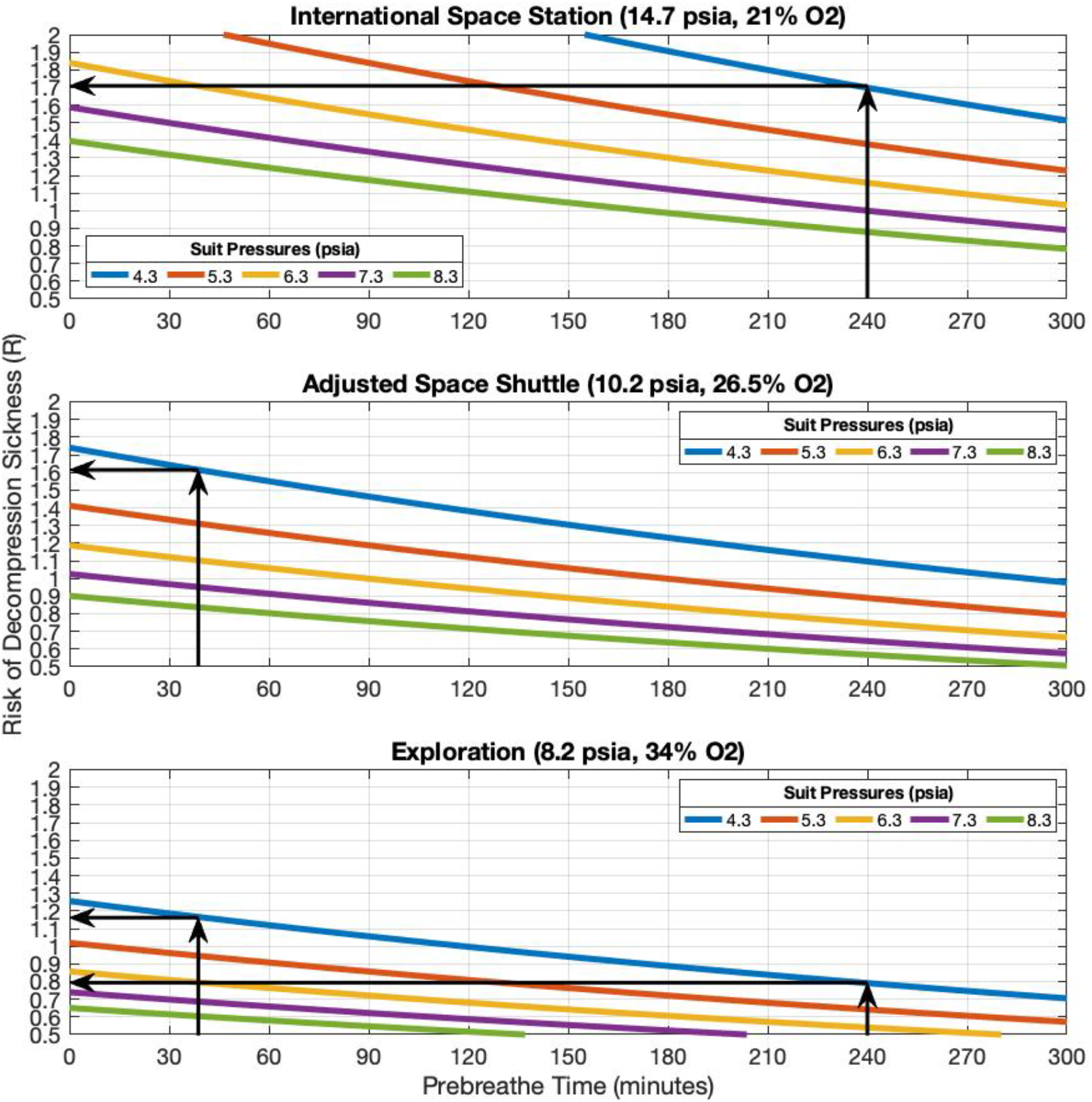
The risk of decompression sickness (DCS) as a function of prebreathe time for multiple spacesuit pressures in ISS (Top Panel), Adjusted Space Shuttle (Middle Panel), and Exploration (Bottom Panel) atmospheric conditions. At each atmospheric condition, higher spacesuit pressures require shorter prebreathe time to maintain a constant DCS risk. In addition, for a constant spacesuit pressure, atmospheric conditions with higher oxygen content (thus less nitrogen) require shorter prebreathe time to maintain a similar DCS risk. For example, in a mission scenario where the objective is to maintain a DCS risk below R = 1.7, a spacesuit at an operating pressure of 4.3 psia (e.g., EMU) requires a prebreathe time of 240 min in the ISS atmospheric conditions, 40 min in the Adjusted Space Shuttle atmospheric conditions, and no prebreathe time in the Exploration atmospheric conditions.

### The Contributions of Mechanical Counterpressure and Gas Pressure to Improved Mobility and Prebreathe Time

There is a clear benefit to increasing the operational spacesuit pressure when attempting to reduce the risk of DCS. However, the increase of gas pressure negatively impacts the mobility of a spacesuit, since gas pressure inhibits joint movement and thus, increases the energy expenditure^51^. In this scenario, the use of MCP can be advantageous, since MCP increases total operating pressure without the negative effects of mobility.

Figure 3 visualizes the trade-off between spacesuit pressure (both gas and MCP), mobility, and risk of DCS in ISS, Adjusted Space Shuttle, and Exploration atmospheric conditions. The background of each panel shows a colored illustration representing the mobility score for a spacesuit given the amount of gas pressure in the spacesuit (x-axis), and we assumed that the mobility score is independent of the amount of pressure that is being provided using MCP. Each panel also includes lines indicating the DCS risk value attained after a one-hour prebreathe time, which is considered to be an acceptable limit for frequent EVA missions in future exploration mission scenarios^22^. For example, in a mission scenario where the habitat conditions are similar to the Adjusted Space Shuttle atmosphere and the spacesuit is purely gas pressurized to 5.3 psi (i.e., no MCP), the allowed 1 hour of prebreathe time reduces the risk of DCS to approximately 1.3, which is in a range recommended by Conkin (1.3-1.4)^22^. In this scenario, the mobility score is ~1.8, which indicates that the energy expenditure for suited ambulation in these conditions is ~1.8 times higher than unsuited walking. If the spacesuit pressure is instead composed of 4.3 psia gas pressure and 1 psia MCP (i.e., total of 5.3 psi), the mobility score decreases to 1.6 (i.e., improved mobility) while maintaining the same DCS risk (R = 1.3). If this same design exercise is conducted using the Exploration atmospheric conditions, we realize that spacesuits with a total pressure of 5.3 psi (either gas pressure alone in combination with up to 2 psia of MCP) present a risk of DCS well below R < 1.3 after one hour of prebreathe time. Finally, in the case of the ISS atmospheric conditions, spacesuits with a total pressure of 5.3 psi (either gas pressure alone in combination with up to 2 psia of MCP) present a risk of DCS well above R > 1.7 after one hour of prebreathe time (indeed, Figure 2, top panel, shows that, if the spacesuit is pressurized to 5.3 psia, the risk of DCS after a 60-min pure oxygen prebreathe becomes R = ~1.95). In all these examples across different atmospheric conditions, we note that mobility scores remain constant since these scores only depend on the amount of gas pressure present in the spacesuit. Figure 3 shows these and others trade-offs between spacesuit gas pressure, spacesuit MCP pressure, and mobility score, across multiple exploration atmospheres.

**Figure 3:**
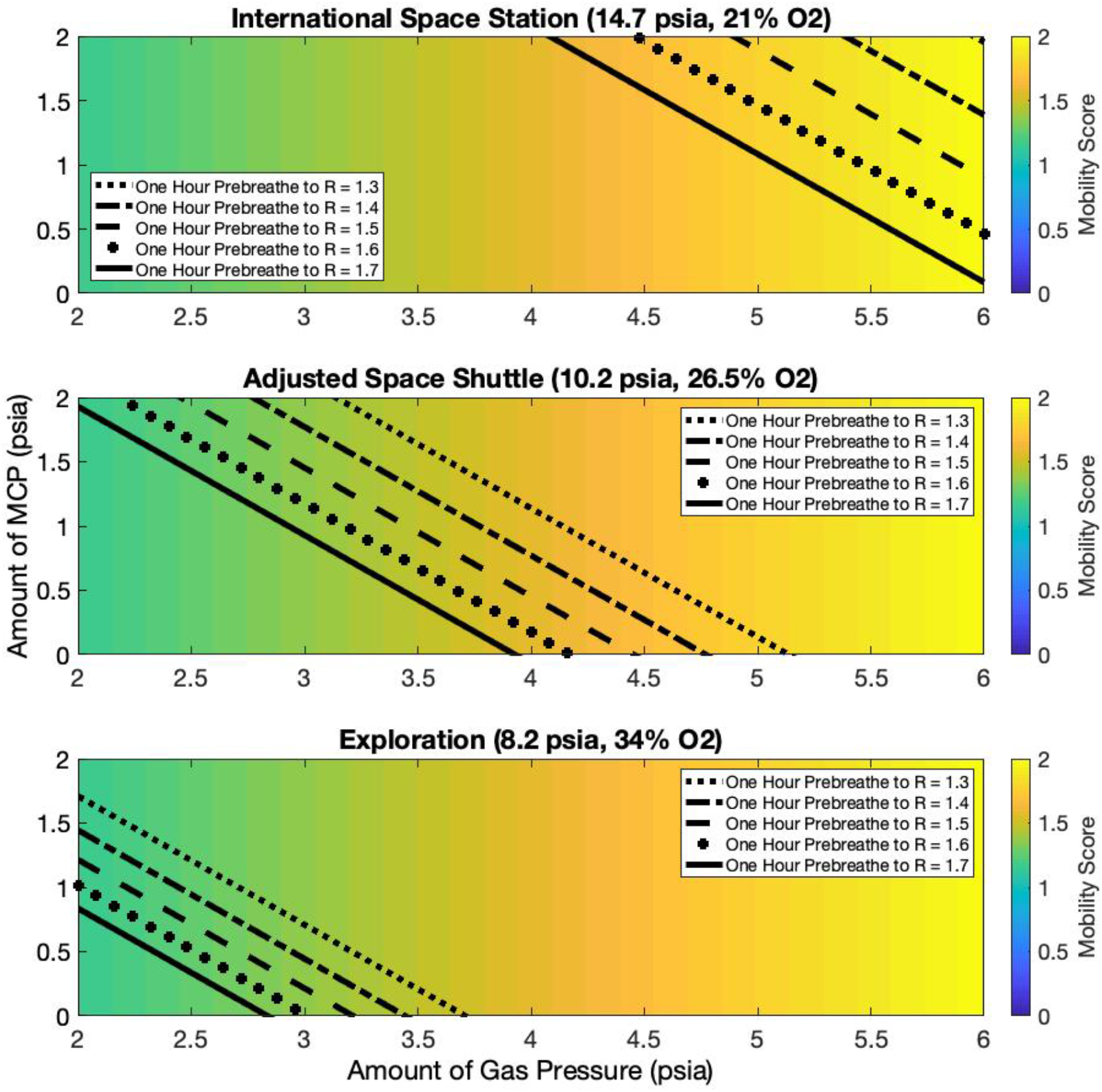
Trade-off between spacesuit pressure, mobility score, and prebreathe times in ISS (Top Panel), adjusted Space Shuttle (Middle Panel), and Exploration (Bottom Panel) atmospheric conditions. The mobility score is a function of gas spacesuit pressure (x-axis) and it is independent to the amount of pressure provided as MCP. One-hour prebreathe times for the given risk score and habitat atmospheric conditions are also indicated in the figures. Mobility improves (i.e., mobility score decreases) with the reduction of gas pressure. At constant prebreathe time (e.g., one hour), the risk of DCS increases with the reduction of total (gas + MCP) spacesuit pressure.

### Benefits of Mechanical Counterpressure in the Spacesuit: Increase in Total Pressure vs. Decrease in Gas Pressure

The addition of MCP to gas pressurized spacesuits, such as the EMU, could result in significant time savings in prebreathe protocols, although these benefits are highly dependent on the atmospheric conditions used in the space habitat. For example, if 1 psia of MCP is added to an EMU-like spacesuit (i.e., 4.3 psia of gas pressure) in the ISS, the prebreathe time needed to attain a DCS risk value of R = 1.7 decreases from 240 min to ~140 min (see Figure 4 top panel and Figure 2 top panel). If 2 psia of MCP are added instead, the prebreathe time needed to attain a DCS risk value of R = 1.7 decreases to ~50 min (see Figure 2 top panel). A similar exercise can be done with other risk values, and other atmospheric conditions, as shown in Figure 4 (which in this case accounts for 1 psia of MCP added to a gas-pressurized spacesuit). We notice that in other atmospheric conditions, such as the Adjusted Space Shuttle and Exploration atmospheres, the operational benefits of adding 1 psia of MCP are not as important. For example, in an Adjusted Space Shuttle atmosphere the traditional EMU gas pressurized spacesuit (i.e., 4.3 psia of gas pressured) only requires 40 min of prebreathe time to attain a DCS risk value of R = ~1.6 (see Figure 4 middle panel). In this scenario, only a small amount of MCP (~ 0.35 psia) will be enough to remove the need for prebreathe activities. However, if the target DCS risk value becomes R = 1.3 (which seems to be more aligned with future planetary exploration requirements), and additional 1 psia MCP decreases the prebreathe time from 153 min to ~44 min. Finally, the addition of MCP to an existing gas-pressurized spacesuit (gas-pressurized to 4.3 psia or higher) in the context of the Exploration atmosphere presents no benefit with respect to prebreathe time, as none is necessary to reach a DCS risk value equal to or lower than R = 1.3 (see Figure 4 bottom panel). Thus, in the Exploration scenario, it is more advantageous to use the MCP capability to reduce the gas-pressure of the spacesuit (as opposed to increase the total pressure), therefore improving mobility.

**Figure 4:**
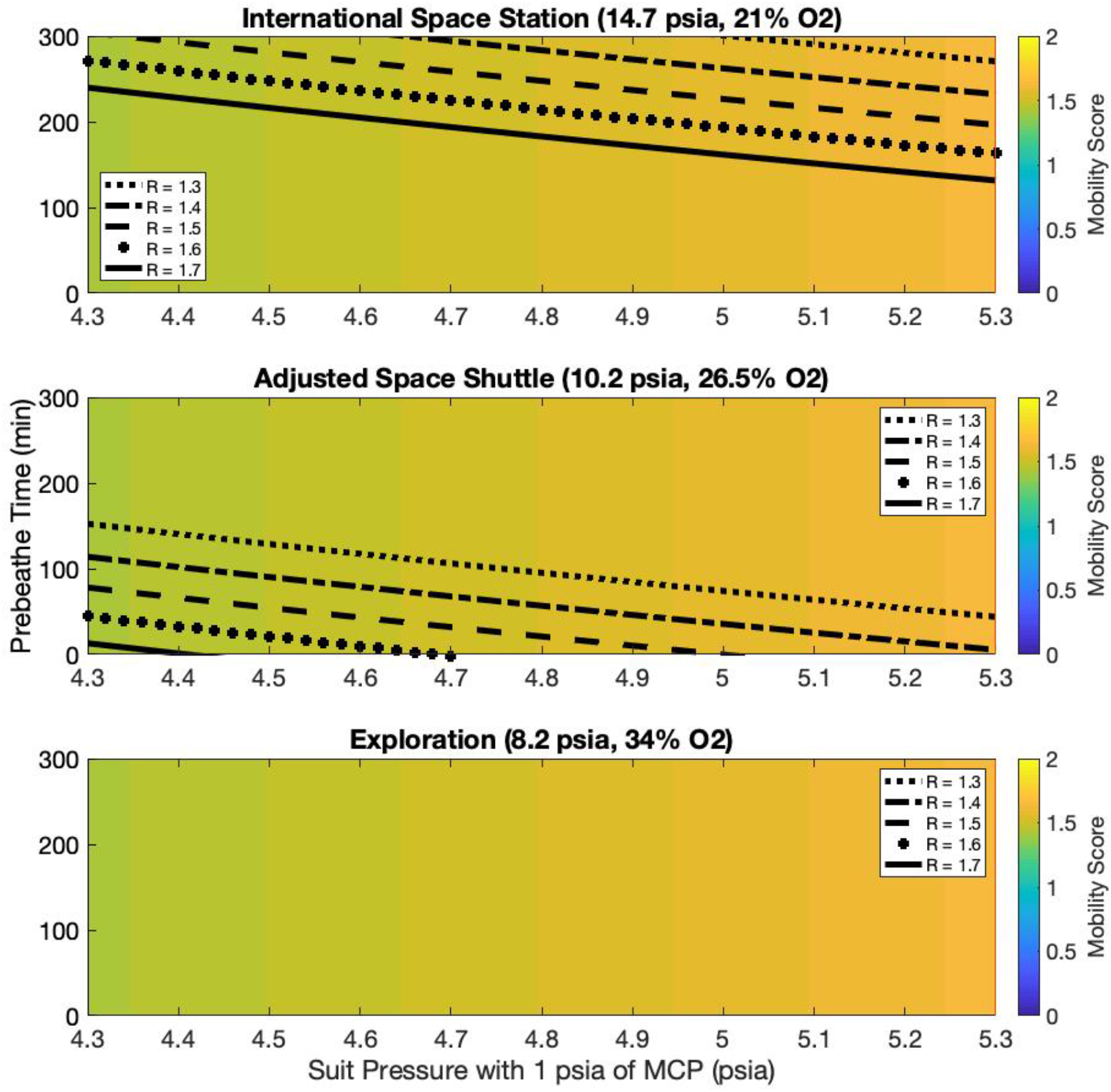
Trade-off between spacesuit pressure (gas pressure plus 1 psia of MCP in x-axis), mobility score, and prebreathe time (y-axis) to attain a given DCS risk value R between 1.7 (currently used in the ISS) and 1.3 (proposed in future planetary exploration) in ISS (Top Panel), adjusted Space Shuttle (Middle Panel), and Exploration (Bottom Panel) atmospheric conditions. The mobility score is a function of gas spacesuit pressure only and it is independent to the amount of pressure provided as MCP. In ISS atmospheric conditions, additional MCP (thus increasing total spacesuit pressure) provides significant operational benefits in reducing prebreathe times. Conversely, in Exploration atmospheric conditions, additional MCP provides no benefits with respect to prebreathing time, as none is necessary to reach an R = 1.3. Thus, in this case it is more advantageous to use MCP to reduce gas pressure (maintaining a constant total pressure) to improve mobility instead.

### Variable Pressure Spacesuit

As the gas pressure of the spacesuit increases, the risk of DCS decreases but mobility becomes more problematic. Conversely, if the gas pressure of the spacesuit decreases, mobility improves, but in order to maintain a similar risk of DCS, either more MCP or a greater prebreathe time becomes necessary. The proposed SmartSuit architecture (i.e., 4.3 psia of gas pressure plus 1 additional psia of MCP), could capitalize on the mobility of a gas-pressurized spacesuit with 4.3 psia, like the EMU, but with a reduced prebreathe time due to a reduced risk of DCS. Additionally, over time and already during the EVA, the SmartSuit architecture allows for a decrease in gas-pressure (e.g., 3.3 psia of gas pressure plus 1 psia of MCP), further increasing mobility. An example of a variable pressure suit is shown in Figure 5. Using for example an Adjusted Space Shuttle atmosphere, a risk value of R = 1.3 can be obtained by completing a prebreathe time of approximately 40 minutes for a suit environment with 4.3 psia of gas pressure and 1 psia of MCP (i.e., total pressure of 5.3 psia). Once a DCS risk value of R = 1.3 is achieved, EVA activities may begin. During the initial EVA activities, the gas pressure of the spacesuit can be steadily decreased from 4.3 to 3.3 psia as the astronauts keep breathing pure oxygen while still sustaining a DCS risk value of R = 1.3. After approximately 150 minutes since the start of the prebreathe activities (as can be found in Figure 2), the spacesuit reaches a total operating pressure of 4.3 psia (3.3 psia gas pressure and 1 psia MCP), which is used for the remainder of the EVA. In this scenario, the mobility score improves from approximately 1.65 at the start of the EVA to just under 1.45 after 150 min of breathing pure oxygen. This framework demonstrates and quantifies the physiological and operational benefits of variable pressure suits, and provides a specific example in the context of the SmartSuit spacesuit architecture. Other examples using other variable pressure spacesuits architectures, such as the xEMU^21^, can easily be incorporated once more details about future EVA planetary operations become available.

**Figure 5:**
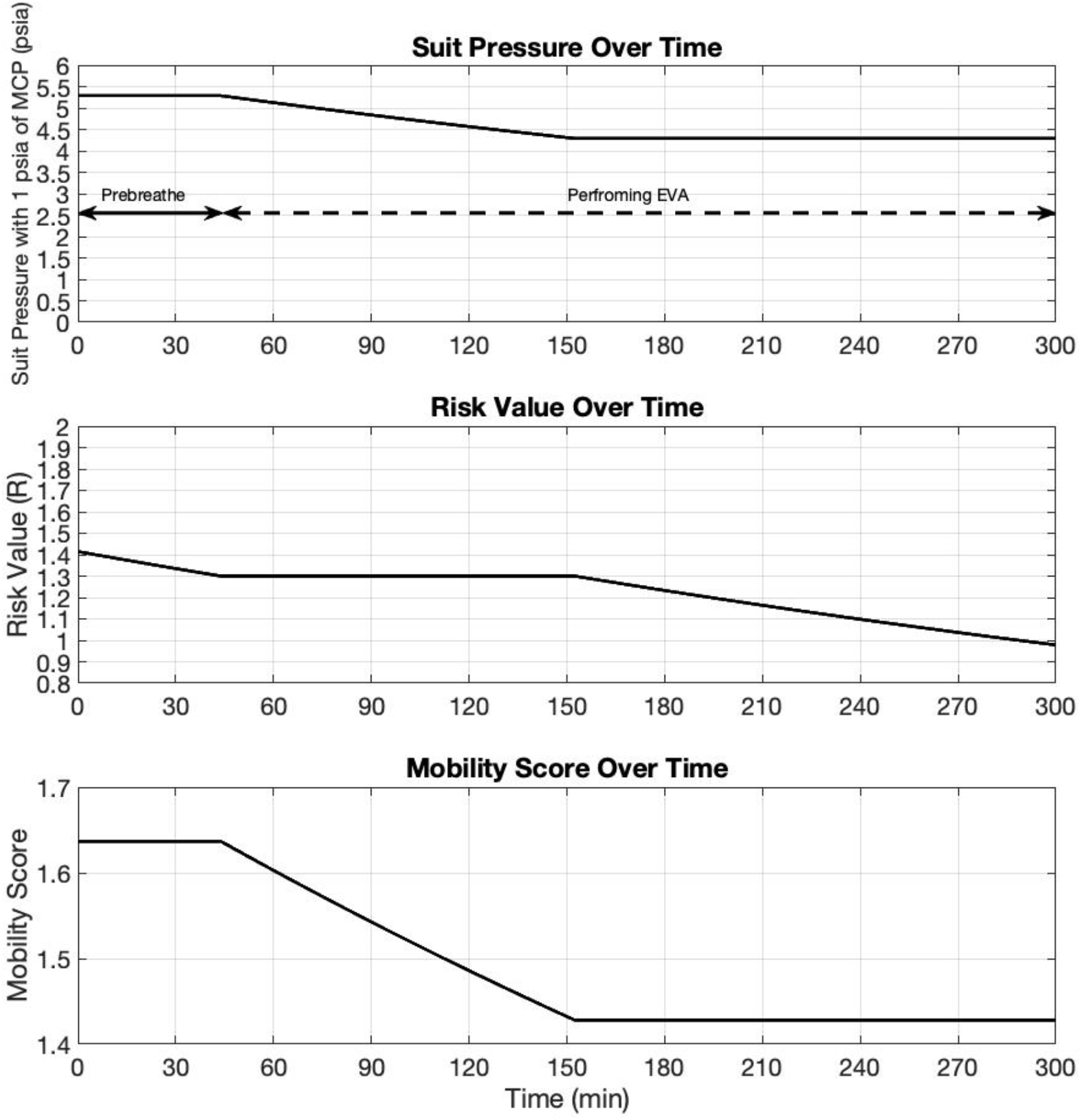
Framework quantifying the physiological and operational benefits of variable pressure spacesuits in the context of the SmartSuit spacesuit architecture, which includes an initial spacesuit pressure of 5.3 psia (4.3 psia of gas pressure and 1 psia of MCP), and a DCS risk value before EVA of R= 1.3 in a habitat with an Adjusted Space Shuttle atmospheric conditions. Prebreathe activities are necessary for 40 minutes to attain a DCS risk value of R = 1.3. Then, the EVA begins while the gas pressure of the spacesuit is steadily decreased as astronauts continue to breath pure oxygen within the spacesuit (while maintaining a DCS risk of R = 1.3). After the 150-minute mark, the spacesuit total pressure of 4.3 psia (3.3 psia of gas pressure plus 1 psia of MCP) is sustained for the rest of the EVA. The reduction of gas pressure from 4.3 psia to 3.3 psia improves the mobility score from approximately 1.65 to just under 1.45.

## Conclusions

We have revisited important trade-offs between spacesuit pressure (both gas pressure and MCP), habitat atmospheric conditions, risk of DCS, and spacesuit mobility in the context of future planetary EVAs. Elevated spacesuit pressures decrease the risk of DCS, but these configurations can have detrimental effects on mobility, which could impact human performance, cause injury, and thus, impact mission success. The use of hybrid spacesuits, which incorporate some level of MCP, are promising in two fronts. First, if the MCP capability is used in *addition to* the gas pressure, this results in a higher total spacesuit pressure, decreasing the risk of DCS (or prebreath time) without major impacts on mobility. On the other hand, if the MCP capability is used to replace part of the gas pressure, this results in a more mobile spacesuit configuration without compromising on DCS risk. We have also demonstrated that these benefits and trade-offs are highly dependent on the atmospheric conditions of the habitat or space station, and therefore, these conditions are also important considerations for future planetary exploration activities.

Finally, we have provided an example of the concept of operations of the SmartSuit, a hybrid spacesuit with a soft robotic layer that, not only increases mobility with assistive actuators in the lower body, but it also applies 1 psia of MCP. The SmartSuit increased mobility encourages a higher operating spacesuit pressure that reduces the risk of DCS, or allow for an increase in cabin pressure, which in turn allows for a lower percentage of oxygen and risk of flammability. The resultant MCP layer can then be used to either increase overall spacesuit pressure (thus, decreasing the risk of DCS or prebreathe time), or to decrease the gas-pressure in the spacesuit (thus, further increasing mobility). These two MCP applications can be effectively combined in the same EVA to maximize the benefits of both configurations, as shown in our example of variable pressure spacesuit.

In conclusion, our framework presented herein quantifies critical trade-offs related to future planetary EVAs activities, contributing to the assessment of human performance during EVAs, and informing spacesuit designers, EVA operation teams, space engineers, and other relevant stakeholders.

## Acknowledgements

This work was supported by the NASA Innovative Advanced Concepts (NIAC) program (grant number 80NSSC19K0969).

## Author Contributions

L.K. and A.D-A conducted the literature review, identified relevant papers, and contributed to written sections of the manuscript. All authors reviewed, edited, and approved the final manuscript.

## Data Availability Statement

All data is open source and available to the public.

